# Recruitment of occipital cortex by arithmetic processing follows computational bias in early blind

**DOI:** 10.1101/319343

**Authors:** Virginie Crollen, Latifa Lazzouni, Antoine Bellemare, Mohamed Rezk, Franco Lepore, Marie-Pascale Noël, Xavier Seron, Olivier Collignon

## Abstract

Arithmetic reasoning activates the occipital cortex of early blind people (EB). This activation of visual areas may reflect functional flexibility or the intrinsic computational role of specific occipital regions. We contrasted these competing hypotheses by characterizing the brain activity of EB and sighted participants while performing subtraction, multiplication and a control verbal task. In both groups, subtraction selectively activated a bilateral dorsal network commonly activated during spatial processing. Multiplication triggered more activity in temporal regions thought to participate in memory retrieval. No between-group difference was observed for the multiplication task whereas subtraction induced enhanced activity in the right dorsal occipital cortex of the blind individuals only. As this area overlaps and exhibits increased functional connectivity with regions showing selective tuning to auditory spatial processing, our results suggest that the recruitment of occipital regions during high-level cognition in the blind actually relates to the intrinsic computational role of the reorganized regions.

## Introduction

Studies involving early blind people (EB) provide important insights into the role played by experience and intrinsic biological constraints in shaping the development of the sensory and cognitive tuning of brain regions. In early blind individuals, regions of the occipital cortex that typically process visual information massively enhance their response to non-visual inputs – a phenomenon called cross-modal plasticity (Bavelier and Neville 2002; Sadato et al. 1996). But what are the mechanisms guiding this cross-modal reorganization process?

It was suggested that these neuroplastic changes are constrained by the native functional organization of the occipital cortex. For example, hMT+/V5, a region chiefly dedicated to processing visual motion in the sighted reorganizes in the blind to preferentially process auditory (Dormal et al. 2016; Jiang et al. 2016; Poirier et al. 2006) and tactile motion (Ricciardi et al. 2007). Moreover, right dorsal occipital regions typically involved in visuospatial localization in the sighted are active when blind individuals localize sounds (Collignon et al. 2011) and altering the function of this region with transcranial magnetic stimulation (TMS) selectively disrupts auditory localization in the blind (Collignon et al. 2007). Similarly, the visual word form area (VWFA), a region specialized to process visual orthographic information in the sighted seems to be selectively recruited in the blind when processing braille words (Büchel et al. 1998; Reich et al. 2011). Those studies suggest that even if the occipital cortex of blind individuals extends their tuning toward non-visual inputs, this reorganization process is not stochastic but is rather constrained by the maintenance of intrinsic computational bias of local regions (Collignon et al. 2009, 2012; Ricciardi et al. 2014; Heimler et al. 2015).

Contrasting with this view, it has been suggested that the occipital cortex of EB engages in higher-level cognitive operations that have apparently little to do with occipital functions such as memory (Amedi et al. 2003), language processing (Bedny et al. 2011; Röder et al. 2002) or numerical thinking (Kanjlia et al. 2016). Based on those observations, it has been proposed that the human cortex is functionally flexible early in life (Lane et al. 2015) and can adopt a wide range of distant computation depending on experience (Bedny 2017).

However, this later argument resides on the presupposition that higher-cognitive functions have no computational relation with vision. But is it the case? Actually, several studies have suggested that the foundations of numerical thinking were rooted in general visuo-spatial mechanisms (Burr and Ross 2008; Ross and Burr 2010; Simon 1999; Stoianov and Zorzi 2012). Arithmetic has for example been thought to involve shifts of attention along a mental number line: a shift of attention toward the right (or toward larger numbers) for addition and toward the left (smaller numbers) for subtraction (Knops et al. 2009a, 2009b; Masson et al. 2014; McCrink et al. 2007; Pinhas and Fischer 2008). Neuroimaging studies have similarly shown that the underlying neural architecture of number representations closely overlaps the one of visuo-spatial processing (Harvey et al. 2013; Shum et al. 2013; Sathian et al. 1999). For instance, topographic numerosity map (numerotopy) in which neural numerosity preferences progress gradually across the cortical surface (Harvey et al. 2013), analogous to sensory maps, have been found in occipital regions typically supporting visuo-spatial/motion processing (Harvey et al. 2017).

The present study was designed to test whether the recruitment of the occipital cortex in early blind individuals by higher-level cognitive functions depends on the intrinsic computational role of specific regions. The study of arithmetic processing represents an interesting test-bed to help disentangling the hypotheses regarding the mechanisms governing cross-modal plasticity (functional recycling versus pluripotency). Indeed, separate arithmetic operations rely on separate brain networks depending on the computational principles they rely on. The resolution of subtraction principally engages a network of dorsal parieto-frontal regions (Chochon et al. 1999; Piazza et al. 2007) presumably due to the "spatial" strategies used to solve such operation (Siegler and Shrager 1984). Like subtraction, multiplication requires the mental manipulation of symbolic numbers, yet it is believed to rely on a distinct temporo-parietal network (Chochon et al. 1999; Zhou et al. 2007) presumably because multiplications are solved by direct fact retrieval (Cooney et al. 1988). This raises the question of whether these different operations produce equivalent neural responses in the occipital cortex of blind individuals, or if, as we presume, subtraction will find a privileged neuronal niche in dorsal occipital regions since these regions keep a privileged role in processing spatial relationship in congenitally blind (Dormal et al. 2012).

We characterized the brain activity of 14 congenitally blind and 16 sighted participants while verifying the results of subtractions and multiplications. If functional reorganization is similarly observed in congenitally blind for both arithmetic operations, this would support the idea that occipital regions are functionally flexible during development and can adopt a wide range of computation depending on experience (Bedny 2017). Alternatively, if solving subtraction problems (which in contrast to multiplication relies on spatial strategies) specifically engages regions of the dorsal occipital cortex typically involved in visuo-spatial processing, this will support the idea that the take-over of occipital regions by higher cognitive functions in the blind actually relies on the original computation of the reorganized regions.

## Results

### Behavioral results

Participants' performances were analyzed with a 3 (experimental conditions: subtraction, multiplication, letter) × 2 (groups: SC vs. CB) repeated measures ANOVA performed on the percentage of correct responses. This analysis showed a marginal effect of condition,++ *F*(2, 56) = 3.13, *p* = .05, 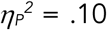. Both groups tended to be less efficient in the subtraction condition (*M* ± *SE* = 87.07 ± 2.55) than in the 2 other conditions (*M* ± *SE* = 91.38 ± 1.29 for multiplication; *M* ± *SE =* 91.97 ± 1.80 for letters). The group effect was not significant, *F*(1, 28) = 0.09, *p* = .76, 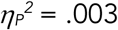, nor was the group x condition interaction, *F*(2, 56) = 0.76, *p* = .47, 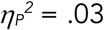 (see Figure 1j).

**Figure 1.**
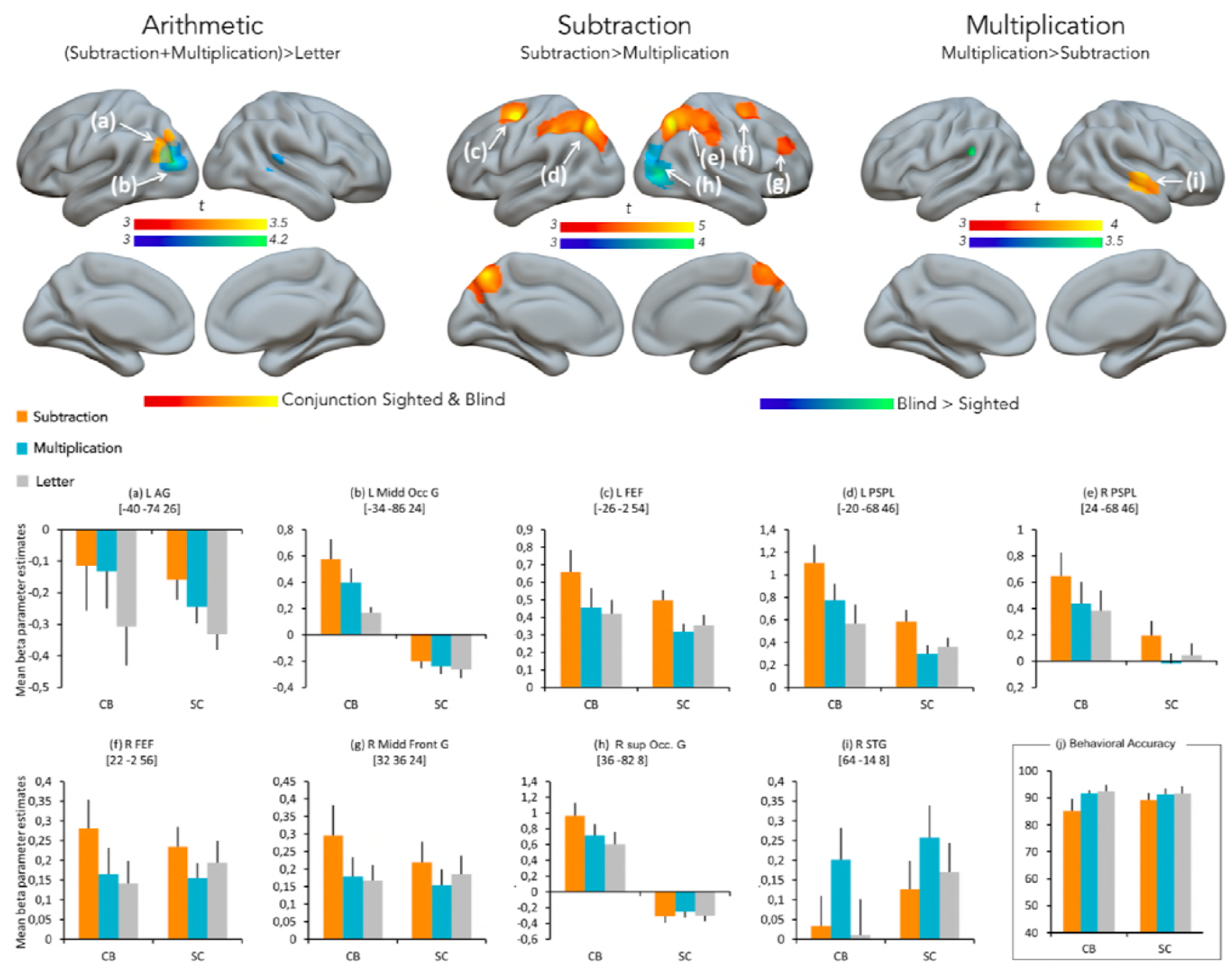
Results of the whole brain univariate analyses. Task-related brain activity common to both groups (puncorr < .002, k > 15) and specific to the blind group (puncorr < .001, k > 15). There were no activations specific to the sighted group. Color bars represent t-values. Lower part (a-i): for illustration, mean activity estimates (arbitrary units ± SEM) associated with arithmetic, subtraction and multiplication are plotted for blind and sighted at significant peaks. See Table 1 for a list of brain regions depicted in this figure. (j) Behavioral results. Mean and standard error of accuracy scores (percentage of correct responses) per condition and group.

### fMRI results

#### General Arithmetic

A conjunction analysis performed across groups disclosed arithmetic selectivity in the left Angular Gyrus (AG) for both groups. To investigate the effect of congenital blindness on global arithmetic processing, we compared the cerebral responses of blind vs. sighted participants for both multiplication and subtraction relative to the letter condition ([CB > SC] [subtraction ∩ multiplication > letter]). This analysis yielded significant results in the left middle occipital gyrus (see Figure 1 and Table 1). The opposite contrast ([SC > CB] [subtraction ∩ multiplication > letter]) did not yield any significant effect.

**Table 1.**
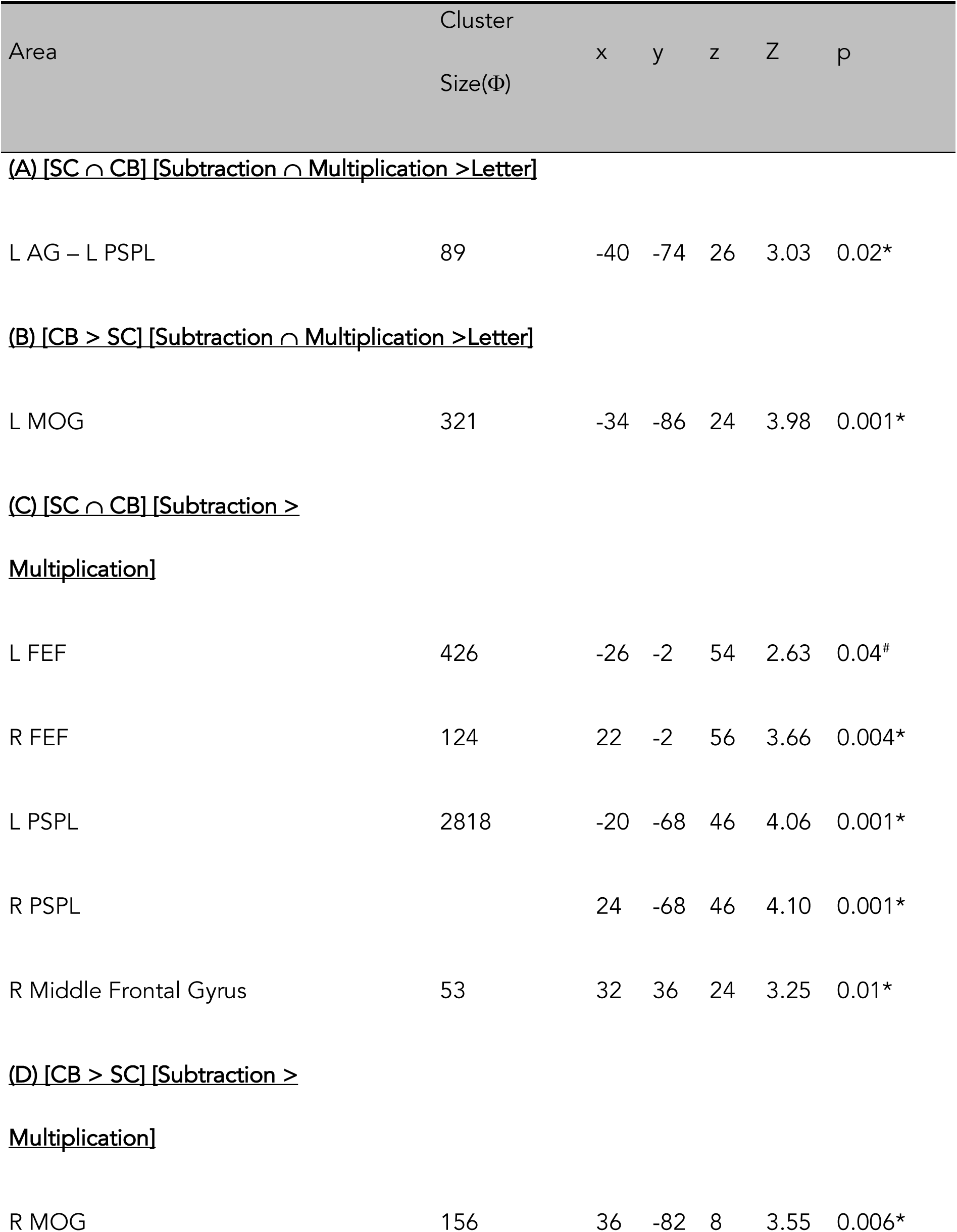
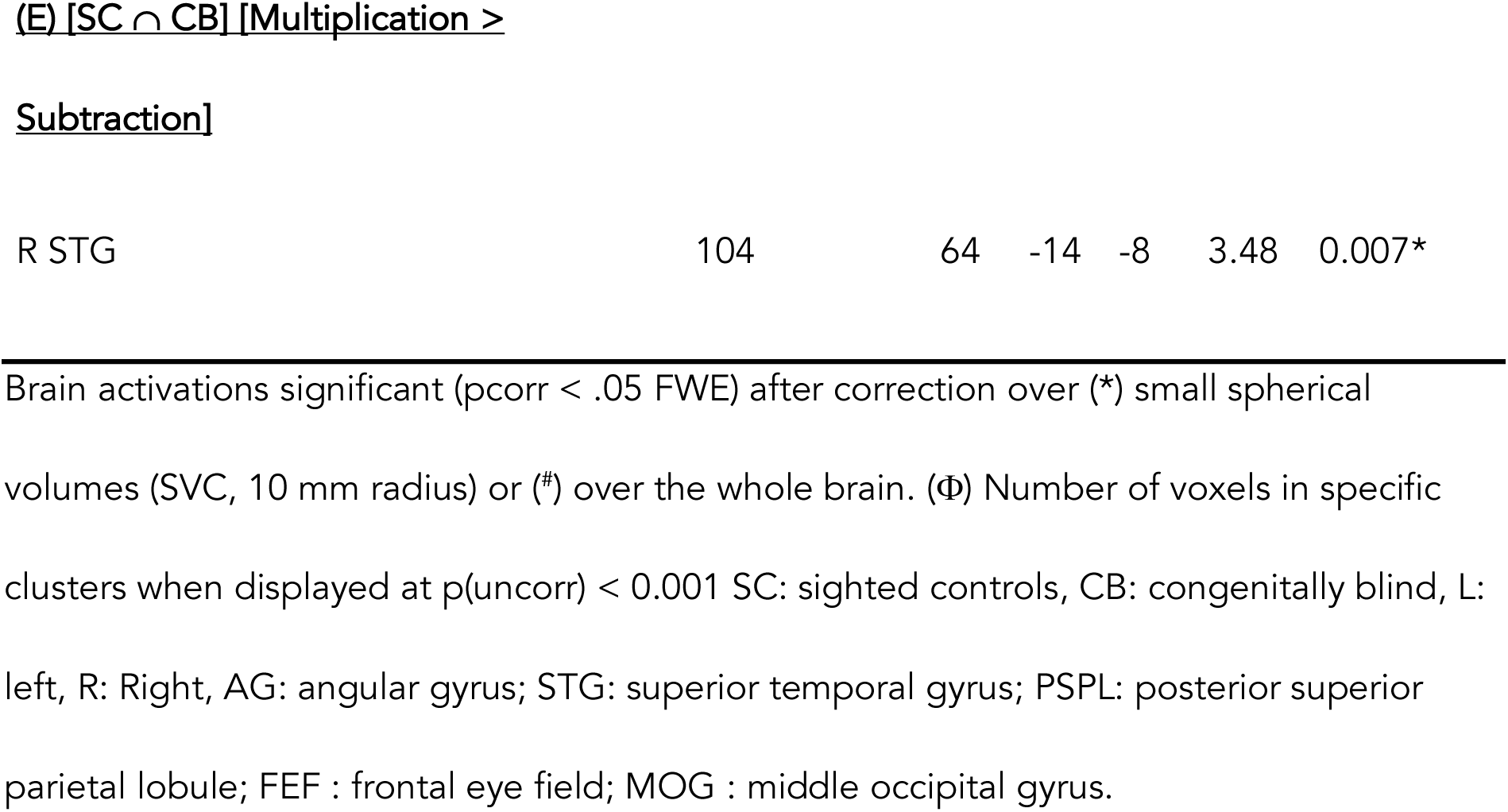
Functional results summarizing the main effect of groups for the different contrasts of interests

#### Subtraction

The frontal eye field (FEF) and the PSPL demonstrated enhanced activity for subtraction over multiplication in both groups of participants ([SC ∩ CB] [subtraction > multiplication]). Crucially, the right middle/superior occipital gyrus (MOG) demonstrated enhanced activity in CB when compared to SC ([CB > SC] [subtraction > multiplication]). In general, deactivation of this region was found in the sighted while activation was observed in CB (see Figure 1 and Table 1). Interestingly and as shown in Figure 2, part of this region also shows selective tuning to auditory spatial processing in the blind (7). The opposite contrast ([SC > CB] [subtraction > multiplication]) did not yield any significant effect.

**Figure 2.**
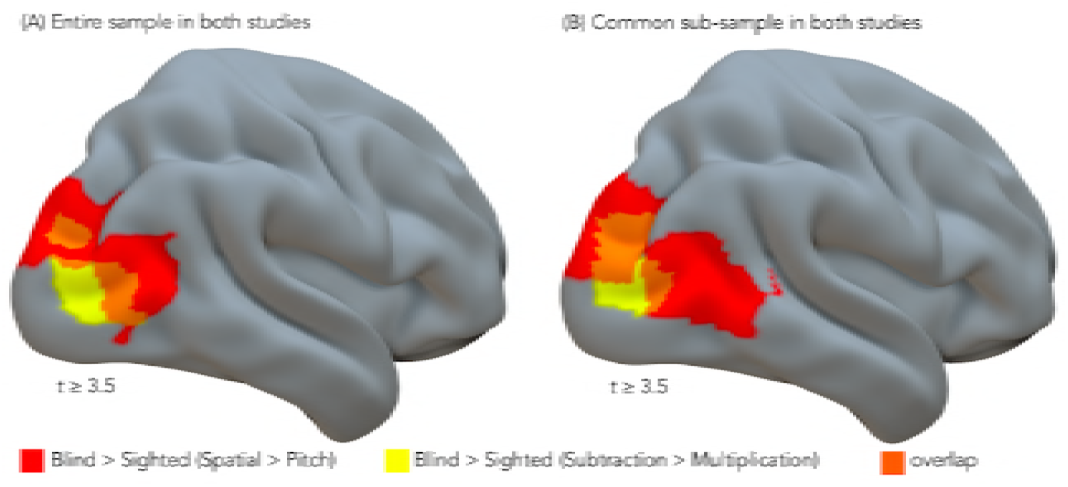
Consistency across studies. The middle occipital gyrus, showing functional preference for subtraction in our study overlapped in part with regions showing selective tuning to auditory spatial processing in the blind (Collignon et al., 2011). Test-consistency either included (A) the entire sample of blind participants in both studies or (B) only the blind participants who performed both studies (N = 7).

#### Multiplication

There was common multiplication related activity in SC and CB ([SC ∩ CB] [multiplication > subtraction]) in the right Superior Temporal Gyrus (STG) (see Figure 1 and Table 1). Neither the contrast ([CB > SC] [multiplication > subtraction]), nor the contrast ([SC > CB] [multiplication > subtraction]) yielded significant effects.

#### Functional connectivity

We found increased correlations between the left MOG and a network of dorsal regions in blind relative to sighted participants (main effect of group, seed to whole-cortex analysis P < 0.05, FDR corrected; Figure 3 and Table 2). A similar pattern was observed for the right MOG (P < 0.05, FDR corrected; Figure 3 and Table 2). Moreover, both seed regions showed enhanced intra-occipital connectivity with ventral occipito-temporal regions as previously shown (Burton et al. 2014; Pelland et al. 2017).

**Table 2.**
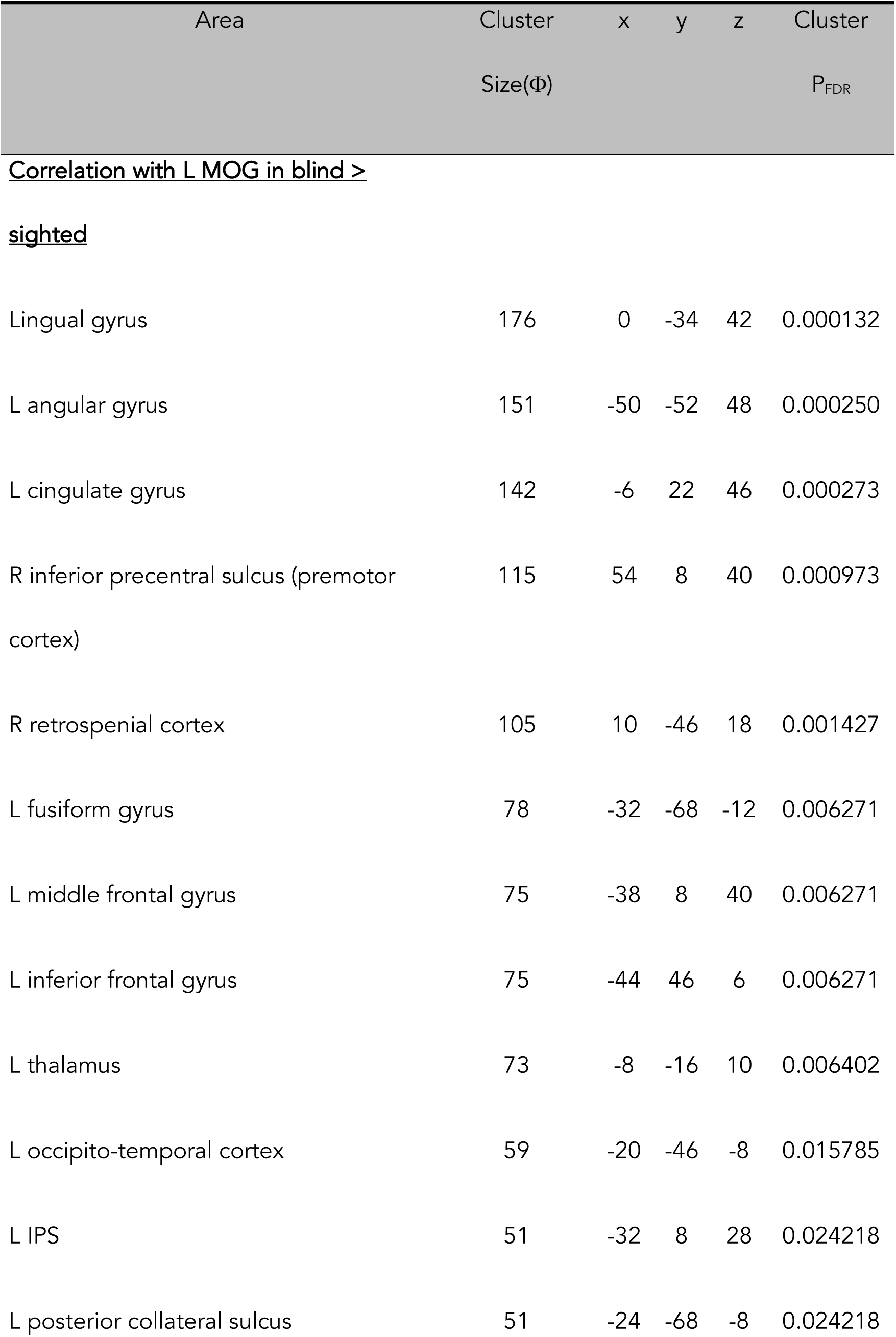
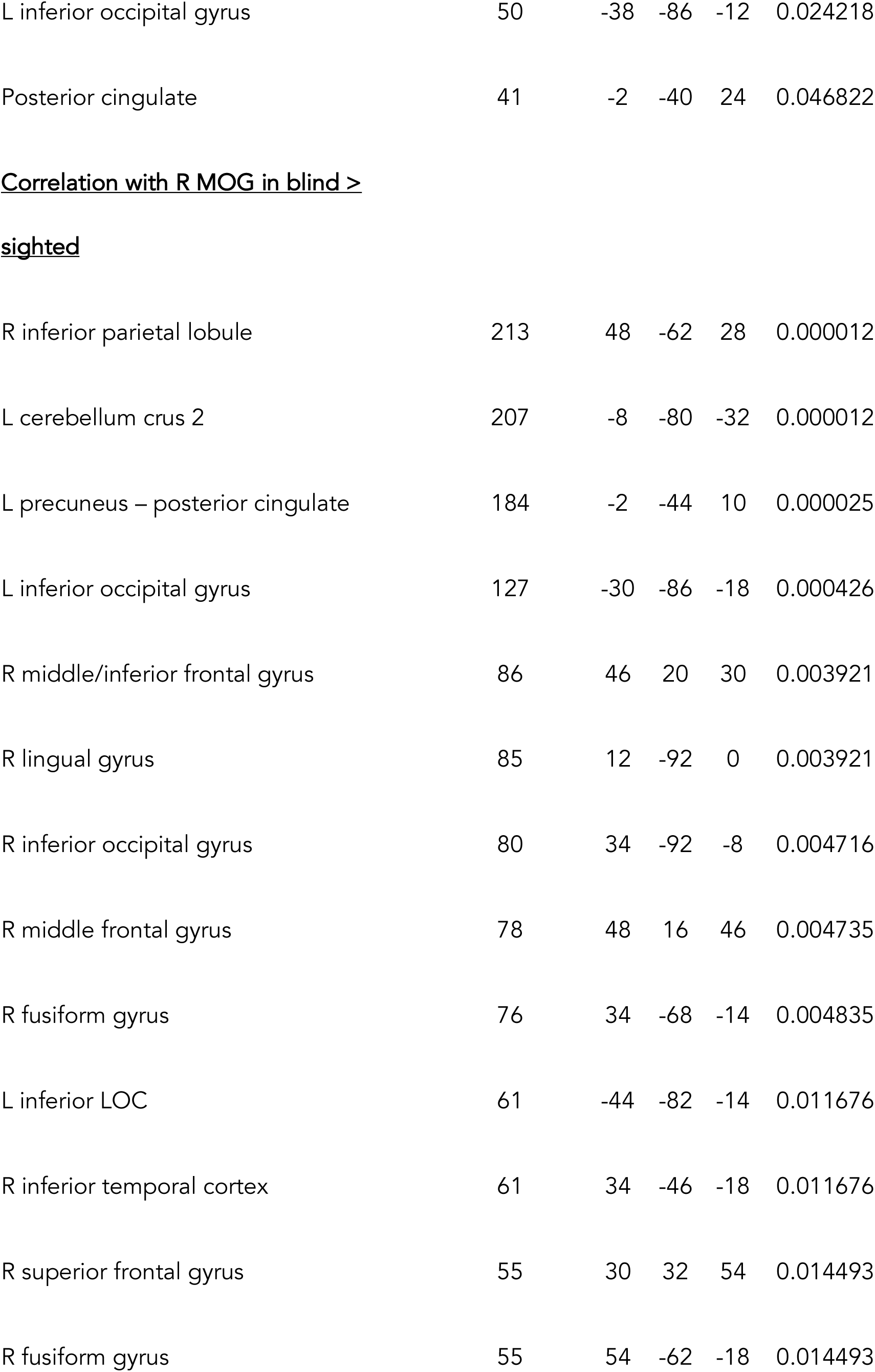
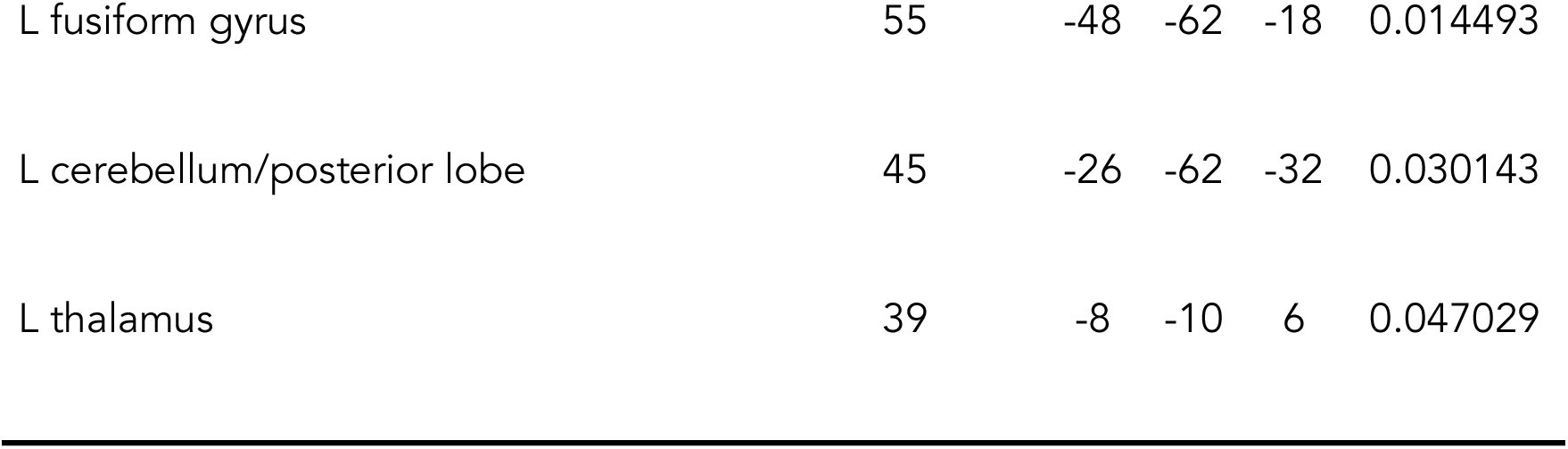
Brain regions correlated with left and right middle occipital gyrus at rest.

| Area | Cluster | x | y | z | Cluster |
| --- | --- | --- | --- | --- | --- |
| | Size($\Phi$) | | | | $P_{FDR}$ |
| <u>Correlation with L MOG in blind ></u> |  |  |  |  |  |
| <u>sighted</u> |  |  |  |  |  |
| Lingual gyrus | 176 | 0 | -34 | 42 | 0.000132 |
| L angular gyrus | 151 | -50 | -52 | 48 | 0.000250 |
| L cingulate gyrus | 142 | -6 | 22 | 46 | 0.000273 |
| R inferior precentral sulcus (premotor<br>cortex) | 115 | 54 | 8 | 40 | 0.000973 |
| R retrosplenial cortex | 105 | 10 | -46 | 18 | 0.001427 |
| L fusiform gyrus | 78 | -32 | -68 | -12 | 0.006271 |
| L middle frontal gyrus | 75 | -38 | 8 | 40 | 0.006271 |
| L inferior frontal gyrus | 75 | -44 | 46 | 6 | 0.006271 |
| L thalamus | 73 | -8 | -16 | 10 | 0.006402 |
| L occipito-temporal cortex | 59 | -20 | -46 | -8 | 0.015785 |
| L IPS | 51 | -32 | 8 | 28 | 0.024218 |
| L posterior collateral sulcus | 51 | -24 | -68 | -8 | 0.024218 |
| L inferior occipital gyrus | 50 | -38 | -86 | -12 | 0.024218 |
| Posterior cingulate | 41 | -2 | -40 | 24 | 0.046822 |
| **Correlation with R MOG in blind > sighted** |  |  |  |  |  |
| R inferior parietal lobule | 213 | 48 | -62 | 28 | 0.000012 |
| L cerebellum crus 2 | 207 | -8 | -80 | -32 | 0.000012 |
| L precuneus – posterior cingulate | 184 | -2 | -44 | 10 | 0.000025 |
| L inferior occipital gyrus | 127 | -30 | -86 | -18 | 0.000426 |
| R middle/inferior frontal gyrus | 86 | 46 | 20 | 30 | 0.003921 |
| R lingual gyrus | 85 | 12 | -92 | 0 | 0.003921 |
| R inferior occipital gyrus | 80 | 34 | -92 | -8 | 0.004716 |
| R middle frontal gyrus | 78 | 48 | 16 | 46 | 0.004735 |
| R fusiform gyrus | 76 | 34 | -68 | -14 | 0.004835 |
| L inferior LOC | 61 | -44 | -82 | -14 | 0.011676 |
| R inferior temporal cortex | 61 | 34 | -46 | -18 | 0.011676 |
| R superior frontal gyrus | 55 | 30 | 32 | 54 | 0.014493 |
| R fusiform gyrus | 55 | 54 | -62 | -18 | 0.014493 |
| L fusiform gyrus | 55 | -48 | -62 | -18 | 0.014493 |
| L cerebellum/posterior lobe | 45 | -26 | -62 | -32 | 0.030143 |
| L thalamus | 39 | -8 | -10 | 6 | 0.047029 |

**Figure 3.**
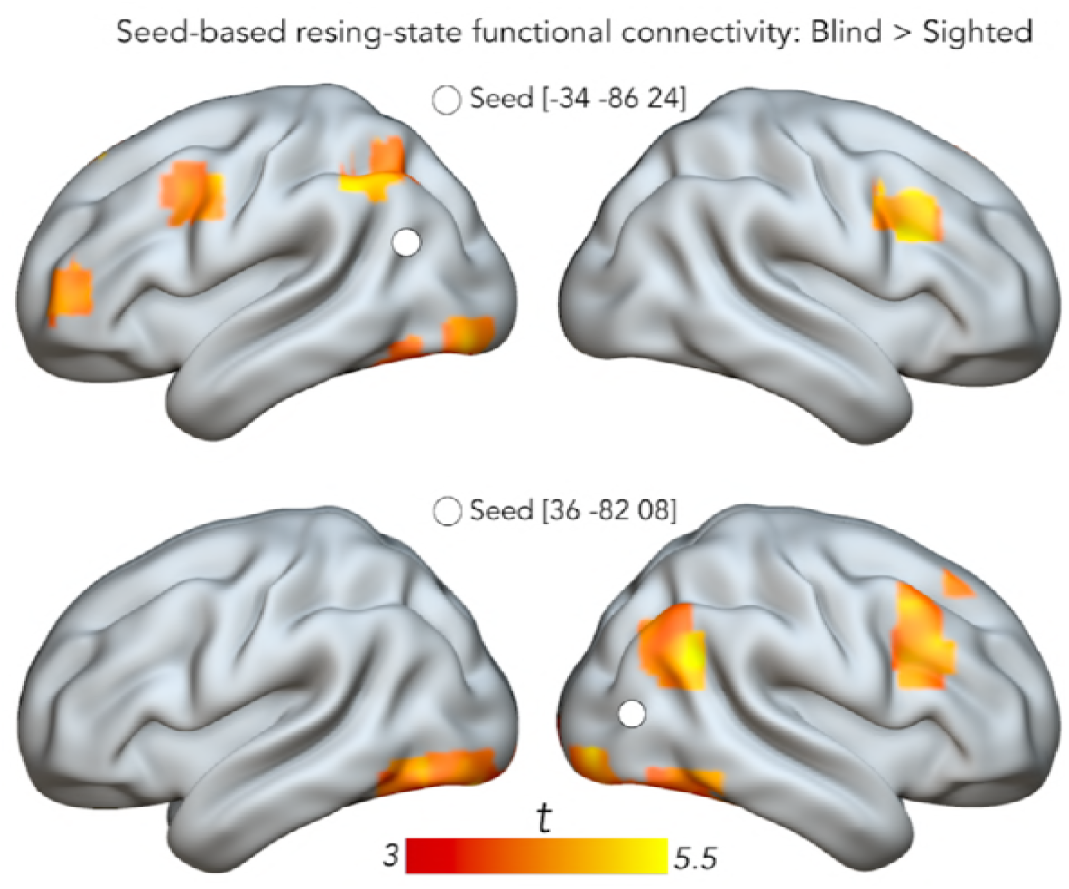
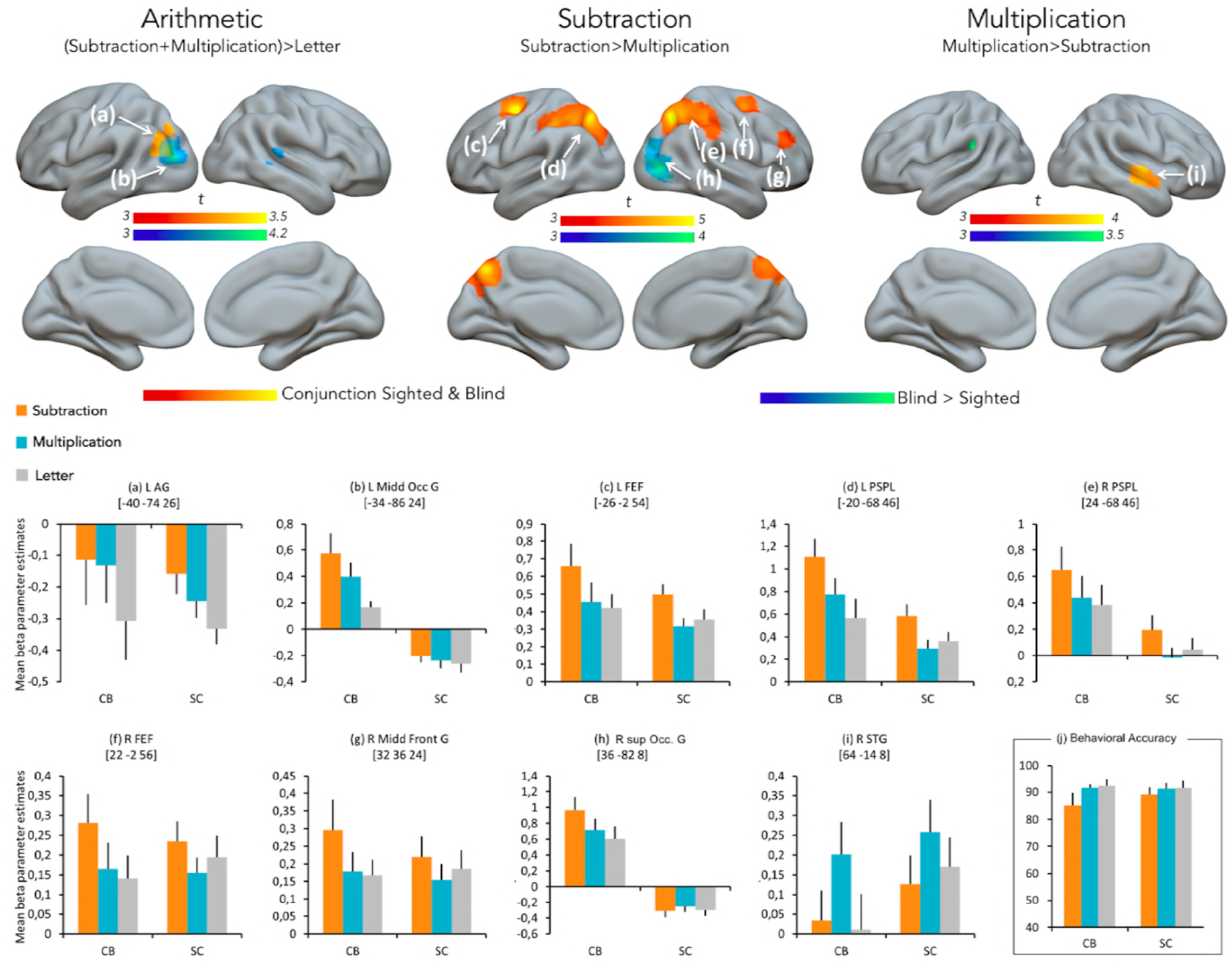
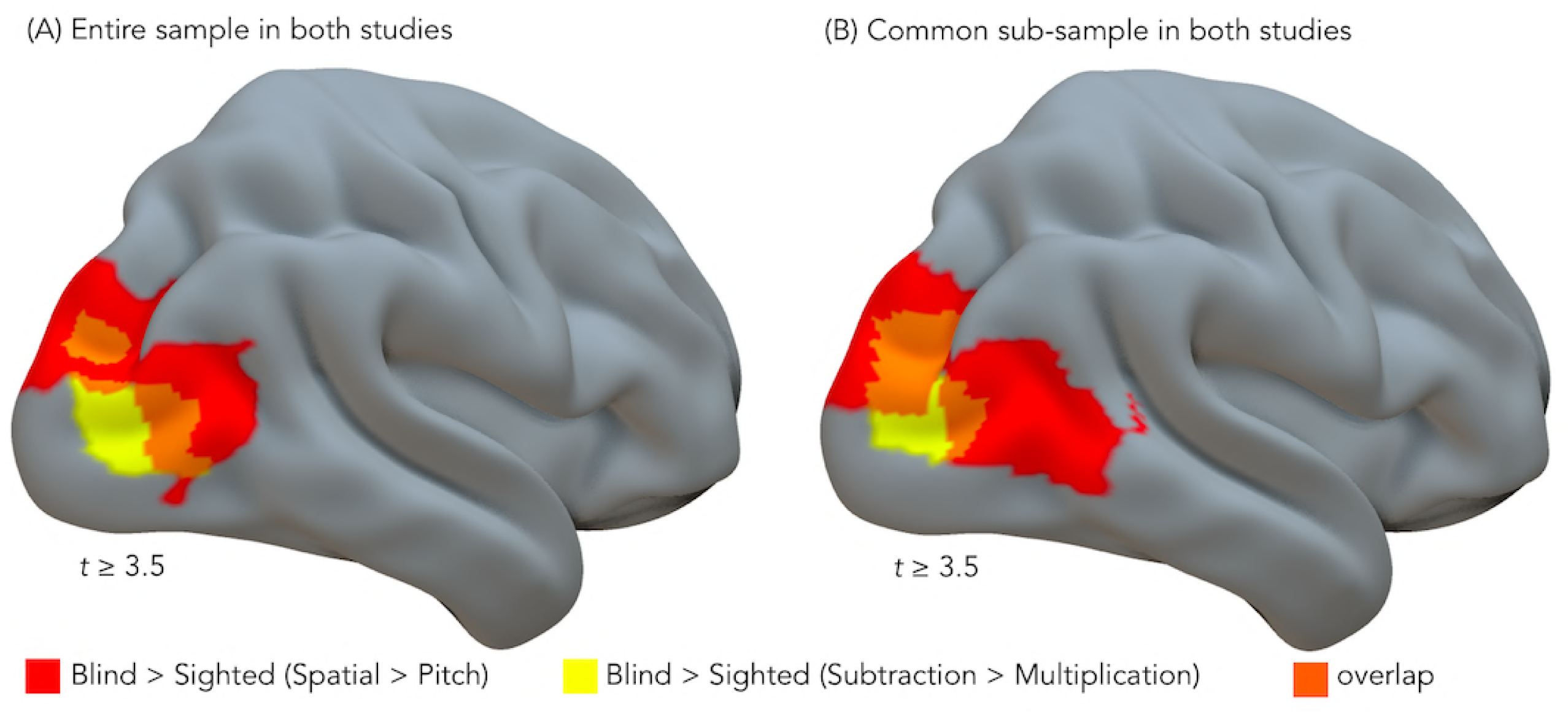
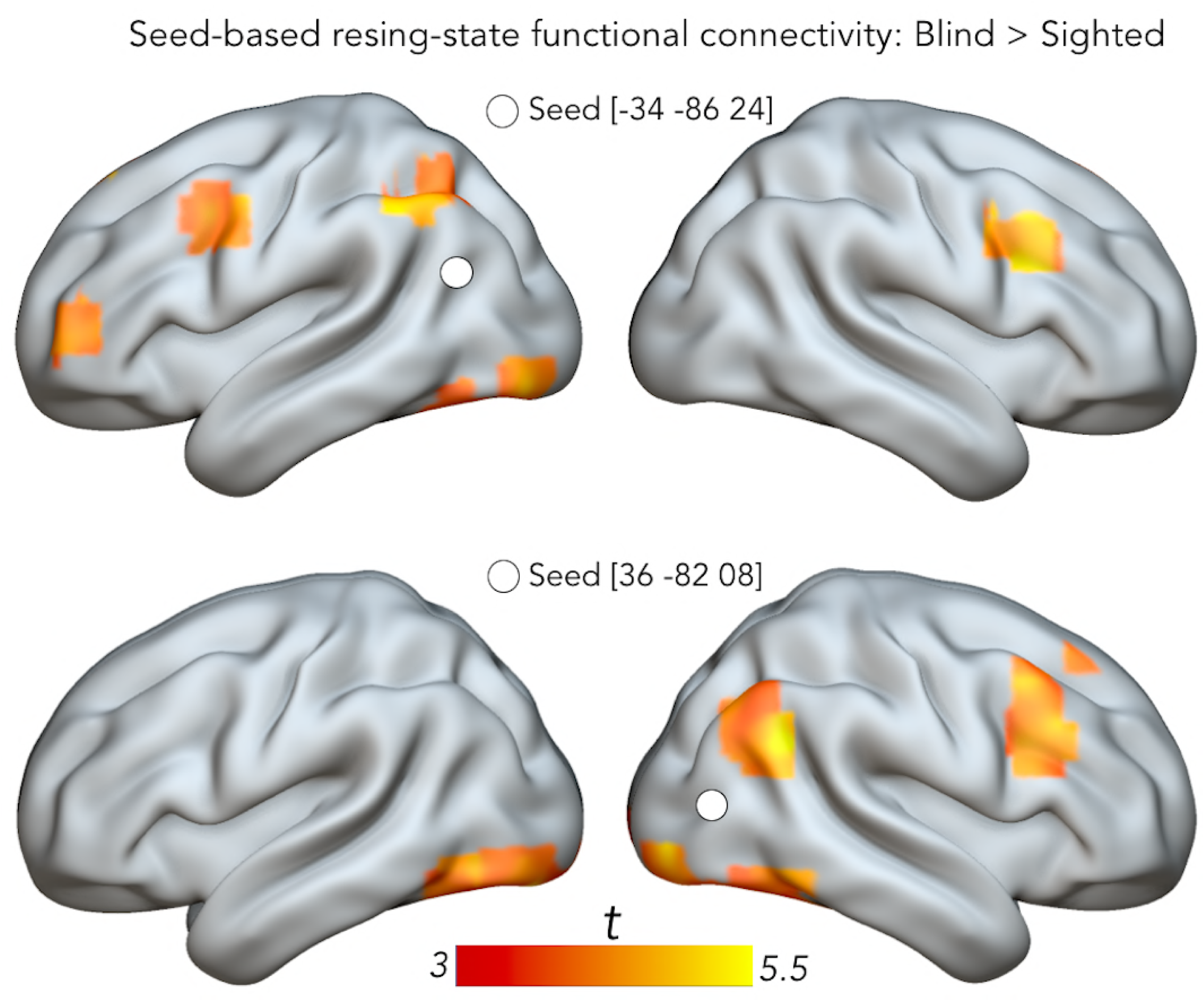
Brain regions more correlated with left and right MOG in blind (n = 13) relative to sighted (n = 11) individuals in resting-state data (P < 0.05, FDR corrected). Left and right MOG seeds shown in white.

## Discussion

We examined how the lack of visual experience impacts on the neuronal basis of specific arithmetical operations by contrasting activity maps elicited by the execution of subtraction or multiplication operations in EB and sighted controls.

Reorganization of occipital regions in case of early blindness provides a unique model for understanding how intrinsic physiology and experience together determine cortical function. Previous studies have found that the "visual" cortex of early blind people responds to a variety of auditory (Collignon et al. 2009; Gougoux et al. 2005; Ricciardi et al. 2014; Weeks et al. 2000) and tactile tasks (Büchel et al. 1998, Sadato et al. 1996).

Several studies have shown that such cross-modal plasticity follows organizational principles that maintain the functional specialization of the colonized brain regions (Amedi, Hofstetter, Maidenbaum, & Heimler, 2017; Bi, Wang, & Caramazza, 2016; Dormal and Collignon, 2011; Cecchetti et al. 2016; Heimler et al. 2015). However, the observation that the occipital cortex of congenitally blind also activates during higher cognitive functions considered distant from visual function, such as arithmetic processing (Kanjlia et al. 2016), was used to challenge the idea that the maintenance of intrinsic computational bias is a generic principle guiding the mechanism of cross-modal plasticity. In contrast, it was supposed that occipital regions are pluripotent early in development and able to engage in a vast array of distant cognitive functions that are evolutionary and cognitively distant from vision (Bedny 2017).

Our observation that specific arithmetic operation – subtraction but not multiplication – triggers enhanced activity in selective occipital regions -the right dorsal stream-provides unifying lights between these two apparently discrepant views of occipital (re)organization in congenitally blind people. Our findings suggest that the cross-modal recruitment of occipital regions for higher cognitive operations does not fully depart from its original function but rather emerges from intrinsic computational bias. More precisely, we suggest that the specific recruitment of dorsal occipital regions for subtraction is a by-product of the intrinsic role of this region for spatial processing (Knops et al. 2009a, 2009b; Masson et al. 2014; McCrink et al. 2007; Pinhas and Fischer 2008). Supporting this idea, region showing selective activation for subtraction in our blind population overlapped in part with regions showing preferential tuning to auditory spatial processing in the blind (Collignon et al. 2011; see Figure 3).

It was proposed that specific numerical processing systematically maps onto parietal circuits because this culturally new invention grounds on, or recycle, more 'basic-primitive' cognitive skills, like space perception or body manipulation, which are evolutionary more ancient (Dehaene and Cohen 2007). We propose that a similar mechanism of functionally specific cortical recycling operates at the ontogenetic level due to experience-dependent neuroplasticity triggered by blindness. Here, subtraction selectively remaps onto dorsal occipital regions due to its reliance on space processing, known to be preserved in these regions in congenitally blind individuals (Dormal et al. 2012). We therefore propose that the recruitment of occipital regions in the blind, even for higher cognitive abilities, finds its "neuronal niche" into a set of circuits that are sufficiently close to the required function and sufficiently plastic as to strengthen or reorient a significant fraction of their neural resources for the non-visual function (Collignon et al. 2009). Interestingly, this raises the possibility that other higher cognitive domains like language, known to also remap in the occipital cortex of the blind (Bedny et al. 2011; Röder et al. 2002), also grounds on the native computational predisposition of these regions (Hasson et al. 2016; van Ackeren et al. 2017).

What could be the mechanistic force guiding this functionally specific reorganization? As illustrated by our connectivity analyses (see Figure 3), the right dorsal occipital region showing enhanced preferential involvement for subtraction in EB also shows enhanced connectivity with a right dorsal network typically involved in spatial processing and attention (Corbetta et al. 1995; Yantis et al. 2003). This result supports the idea that a biased inborn connectivity profile between brain regions may guide the functional specialization of brain regions and, by extension, constraints how cross-modal plasticity expresses in the occipital cortex of the blind (Hannagan et al. 2015). In the context of our study, the reinforcement of a privileged connection between dorsal occipital regions and the intraparietal sulcus, probably rooted on common involvement in spatial computation, will extend the preserved function of the parietal cortex (eg. for subtraction) toward dorsal occipital regions in case of early visual deprivation.

This additional involvement of dorsal occipital regions for subtraction in EB inevitably raises the question of what is happening to the network typically involved in this function. Even if in our study we did not find any enhanced activity in sighted versus blind participants, previous studies have suggested that the enhanced occipital involvement in EB may be concomitant to a reduction of the computational load of the typical network (Dormal et al. 2016).

The left AG was the unique region in which both subtraction and multiplication produced superior activity than the letter control task in both sighted and blind individuals (Figure 1). Precisely, this region was less deactivated during calculation than when processing letters in both groups. Deactivation has been consistently reported during arithmetic tasks (Grabner et al. 2007; Mizuhara et al. 2005; Rickard et al. 2000; Wu et al. 2009; Zhou et al. 2007) and has been shown to be negatively correlated to mathematical performance: stronger is the deactivation, lower is the arithmetic performance (Wu et al. 2009).

More importantly, we demonstrate that specific arithmetic operations activate selective networks: subtraction preferentially activated the FEF and the PSPL regions while the right STG was activated for multiplication in both groups. This dissociation between parieto-frontal regions for subtraction and superior temporal regions for multiplication supports the idea that arithmetic is processed in different formats within distinct cerebral pathways (Dehaene and Cohen 1997). Therefore, visual experience does not have a foundational role in setting-up the functional segregation between subtraction and multiplication. The evidence that the neural representation of numbers is not tied to visual abilities or experience demonstrate that numerical concepts can be acquired through non-visual mechanisms (Dormal et al. 2016; Crollen et al. 2017). A recent study also showed that blind individuals activate parietal regions to solve subtraction (Kanjlia et al. 2016). However, this study did not investigate functional selectivity for specific arithmetical operation. We therefore confirm and extend these results by showing that the dissociation of brain regions supporting specific arithmetic operation is preserved in blind people.

Does this overlap of the brain circuits involved in arithmetic processing in early blind and sighted individuals means that the way these regions implement numerical processing is immune to visual input and experience? One possibility is that those selective neural networks supporting specific numerical operations rely on computational procedures, like memory retrieval for multiplication and spatial processing for subtraction (Campbell and Xue, 2001). These procedures may be abstracted from sensory input and experience and may therefore be built on amodal representational format (Damarla et al. 2016; Eger et al. 2003; Nieder 2012, 2016). However, the fact that similar regions activate in blind and sighted individuals during the same specific task (i.e., parietal network during subtraction) does not guarantee that the specific format of the cognitive operation is similar across both groups. Indeed, if the lack of vision does not preclude the optimal development of various numerical skills (Dormal et al. 2016; Castronovo 2014; Crollen and Collignon 2016), some qualitative properties of numerical representations seems however to critically depend on early visual experience. For example, early blindness changes the nature of the reference frame in which the spatial processing of numbers occurs: while sighted and late blind participants associate numbers to an external frame of reference, congenitally blind individuals principally rely on an association between numbers and an egocentric coordinate system (Crollen et al. 2013). Blindness also alters the typical development of finger-counting, a procedure often used by sighted individuals while learning basic addition and subtraction operations (Crollen et al. 2011, 2014). Given that early blindness affects the use of an external visuo-spatial frame of reference and the implementation of finger-counting, it is possible that the common regions involved in numerical processing in both groups rely on distinct representational format in early blind and sighted individuals. It will therefore be important for future studies to assess whether the representations of the numerical information embedded in those brain circuits are truly independent of visual experience or if visual experience influences the format of these representations despite overlapping activation.

## Materials and Methods

### Participants

Sixteen sighted controls (SC) [6 females, age range 22-64 y, (mean ± SD, 44 ± 14 y)] and 14 congenitally blind (CB) participants [3 females, age range 23-61 y, (mean ± SD, 44 ± 13 y)] took part in the study (see supplemental table 1 for a detailed description of the CB participants). The SC did not statistically differ from the CB group for age (*t*(28) = -0.02, *p* > .9) and sex ratio (*χ*^2^ = 1.54, *p* = .21). The participants in the blind group were totally blind since birth or had, at the utmost, only rudimentary sensitivity for brightness differences and never experienced patterned vision (never saw colors, shapes, or motion). In all cases, blindness was attributed to peripheral deficits with no additional neurological problems. Procedures were approved by the Research Ethics Boards of the University of Montreal. Experiments were undertaken with the understanding and written consent of each participant. Sighted participants were blindfolded when performing the task.

### Task and general experimental design

Participants were scanned in 1 fMRI session using a block design procedure. During scanning, participants had to: 1) verify the result of subtractions; 2) verify the result of multiplications; and 3) perform vowel/consonant judgment verification on letters. Additions were not presented as they are thought to not only involve spatial displacements on the mental number line but are also assumed to rely on rote verbal memory (Dehaene & cohen, 1997). In the subtraction verification task, triplets of auditory numbers were presented and participants had to judge whether the third number corresponded to the difference of the first two numbers. The first operand was either 11 or 13; the second operand ranged from 3 to 8. The third number was either the correct result of the subtraction or the correct result ± 1. In the multiplication verification task, triplets of auditory numbers were presented and participants had to judge whether the third number corresponded to the product of the first 2 numbers. The first operand was either 3 or 4. The second operand ranged from 3 to 8. The third number presented corresponded either to the correct result or to the correct result ± the first operand (e.g., three, five, twelve). In order to examine the neural activity of arithmetic in general (common activity for subtraction and multiplication), participants had also to perform a control letter task. This task was matched to the numerical tasks in terms of stimuli presentation (3 consecutive – non-numerical – symbolic stimuli) and response requirements. Triplets of letters were thus auditory presented and participants had to judge whether the third letter pertained to the same category (vowel vs. consonant) as the first 2 letters (the first 2 letters were always of the same category). The letters used were the vowels A, E, I, O, U and the consonants B, D, M, N, P, R, S. Participants responded with their right index finger using the top (for correct triplet) or the bottom (for incorrect triplet) key of a response box.

The fMRI session consisted of 30 successive blocks (24 s duration each) alternating the three tasks in a fixed order and separated by rest periods ranging from 7 to 9 s (median 8 s). Each block consisted of 6 successive auditory triplets of 4000 ms. In the scanner, auditory stimuli were delivered by means of circumaural fMRI-compatible headphones (Mr Confon, Magdeburg, Germany).

Before the fMRI acquisition, all participants underwent a training session in a mock scanner with recorded scanner noise played in the bore of the stimulator to familiarize them with the fMRI environment and to ensure that the participants understood the task.

### fMRI data acquisition and analyses

Functional MRI-series were acquired using a 3-T TRIO TIM system (Siemens, Erlangen, Germany), equipped with a 12-channel head coil. Multislice T2*-weighted fMRI images were obtained with a gradient echo-planar sequence using axial slice orientation (TR = 2200 ms, TE = 30 ms, FA = 90°, 35 transverse slices, 3.2 mm slice thickness, 0.8 mm inter-slice gap, FoV = 192×192 mm^2^, matrix size = 64×64×35, voxel size = 3×3×3.2 mm^3^). Slices were sequentially acquired along the z-axis in feet-to-head direction. The 4 initial scans were discarded to allow for steady state magnetization. Participants' head was immobilized with the use of foam pads that applied pressure onto the headphones. A structural Tl-weigthed 3D MP-RAGE sequence (voxel size= 1×1×1.2 mm3; matrix size= 240×256; TR= 2300 ms, TE= 2.91 ms, TI= 900 ms, FoV= 256; 160 slices) was also acquired for all participants.

Functional volumes were pre-processed and analyzed using SPM8 (http://www.fil.ion.ucl.ac.uk/spm/software/spm8/; Welcome Department of Imaging Neuroscience, London), implemented in MATLAB (MathWorks). Pre-processing included slice timing correction of the functional time series (Sladky et al. 2011), realignment of functional time series, co-registration of functional and anatomical data, a spatial normalization to an echo planar imaging template conforming to the Montreal Neurological institute space, and a spatial smoothing (Gaussian kernel, 8mm full-width at half-maximum, FWHM).

Following pre-processing steps, the analysis of fMRI data, based on a mixed effects model, was conducted in two serial steps accounting respectively for fixed and random effects. For each subject, changes in brain regional responses were estimated through a general linear model including the responses to the 3 experimental conditions (subtractions, multiplications, letters). These regressors consisted of a boxcar function convolved with a canonical double-gamma hemodynamic response function. Movement parameters derived from realignment of the functional volumes (translations in x, y and z directions and rotations around x, y and z axes) and a constant vector were also included as covariates of no interest. High-pass filtering was implemented in the design matrix using a cut-off period of 128 s to remove slow drifts from the time series. Linear contrasts tested the main effect of each condition ([Subtraction], [Multiplication], [Letter]), the main effect of arithmetic ([Subtraction ∩ Multiplication>Letter]), the main effect of the Subtraction condition ([Subtraction>Multiplication]) and the main effect of the multiplication condition ([Multiplication>Subtraction]). These linear contrasts generated statistical parametric maps [SPM(T)]. The resulting contrast images were then further spatially smoothed (Gaussian kernel 8 mm FWHM) and entered in a second-level analysis, corresponding to a random effects model, accounting for inter-subject variance. For each contrast, one-sample t-tests were carried out in each group separately. Two-sample t-tests were then computed to identify group differences for each separate contrast. Group effects [Blind>Sighted] and [Sighted>Blind] were inclusively masked (p < 0.001 uncorrected for multiple comparisons) by the main effect in the blind and the sighted group, respectively. Statistical inferences were performed at a threshold of p < 0.05 after correction for multiple comparisons (Family Wise Error method) over the entire brain volume or over small spherical volumes (10 mm radius) located in regions of interest (SVC). To select the coordinates of interest, we consulted a body of literature examining brain activations related to numerical processing in sighted individuals and related to functionally specific cross-modal plasticity in the blind. Before performing any small-volume correction (SVC), peaks reported in Talairach space (Talairach and Tournoux 1988) were transformed to Montreal Neurological Institute space using Matthew Brett's bilinear transformation (http://imaging.mrc-cbu.cam.ac.uk/imaging/MniTalairach). Standard stereotactic coordinates (x,y,z) used for SVC are listed in supplemental material (in MNI space).

### Resting-State Functional Connectivity Analysis

To control for possible motion artifacts, a scrubbing approach was implemented using Artifact detection tools (ART; www.nitrc.org/projects/artifact detect) to identify outlier volumes that had a difference in scan-to-scan global intensity more than 9 standard deviations away from the mean global brain signal, or volumes that had more than 2 mm of scan-to-scan composite motion. The outlier time points were used as a first level covariate. Afterwards, the functional volumes were spatially smoothed with a 4-mm full-width half-maximum (FWHM) Gaussian kernel. To rule out the possibility that any connectivity changes could be attributed to motion (Power et al. 2012; Satterthwaite et al. 2012; Van Dijk et al. 2012), we compared the average and maximum motion between groups using a two-sample t-test, and found no significant difference between the 2 groups in the average (*t*(22)=0.35, p=0.73) or the maximum motion (*t*(22)=0.67, p=0.508).

Functional connectivity analysis was performed in CONN functional connectivity toolbox (Whitfield-Gabrieli and Nieto-Castanon 2012); http://www.nitrc.org/projects/conn). A component-based noise correction (Compcorr) (Behzadi et al. 2007) strategy was implemented to control for nuisance effects and physiological noise. Motion parameters and their first derivatives, and outlier time points from ART toolbox were included as first level covariates to remove the variance related to head motion. Linear de-trending and a band-pass filter of 0.008-0.09Hz was applied to the functional data. First level analysis was performed in the CONN framework to investigate functional connectivity changes within-subject. Pearson's correlation coefficient was used as a measure of functional connectivity. A seed-based correlation analysis using a voxel-wise approach was performed. We used the left and right middle occipital gyri as our seed regions because these regions showed cross-modal plasticity for the contrasts [CB>SC][Subtraction∩Multiplication>Letter] and [CB>SC][Subtraction>Multiplication] – see results section.

The time series from each seed region was extracted by averaging the signal from all the voxels in the seed ROI. The time series from each seed was then correlated with each voxel in the rest of the brain. The correlation coefficients were fisher transformed to perform second-level statistical comparison across groups. One-sided two-samples t-test was performed to assess differences in the functional connectivity between groups [CB > SC]. Results were thresholded at p<0.001 at the voxel level and FDR-corrected p<0.05 at the cluster level.

## Acknowledgments

The authors are grateful to Giulia Dormal for her help in implementing the design of this study. This research and the authors were supported by the Canada Research Chair Program (FL), the Canadian Institutes of Health Research (FL), the Belgian National Funds for Scientific Research (OC, MPN), a WBI. World grant (VC), the European Union's Horizon 2020 research and innovation program under the Marie Sklodowska-Curie grant agreement No 700057 (VC) and the 'MADVIS' European Research Council starting grant (OC; ERC-StG 337573).

